# Local processing of visual information in neurites of VGluT3-expressing amacrine cells

**DOI:** 10.1101/127159

**Authors:** Jen-Chun Hsiang, Keith Johnson, Linda Madisen, Hongkui Zeng, Daniel Kerschensteiner

## Abstract

Synaptic inputs to neurons are distributed across extensive neurite arborizations. To what extent arbors process inputs locally or integrate them globally is, for most neurons, unknown. This question is particularly relevant for amacrine cells, a diverse class of retinal interneurons, which receive input and provide output through the same neurites. Here, we used two-photon Ca^2+^ imaging to analyze visual processing in VGluT3-expressing amacrine cells (VG3-ACs), an important component of object motion sensitive circuits in the retina. VG3-AC neurites differed in their preferred stimulus contrast (ON vs. OFF); and ON and OFF responses varied in transience and preferred stimulus size. Contrast preference changed predictably with the laminar position of neurites in the inner plexiform layer. Yet, neurites at all depths were strongly activated by local but not by global image motion. Thus, VG3-AC neurites process visual information locally, exhibiting diverse responses to contrast steps, but uniform object motion selectivity.

## Introduction

Neurons receive most of their synaptic input on large intricately branched dendritic arborizations. Traditionally, distributed inputs were thought to be summed linearly at the cell body (Yuste, 2011). However, recent studies uncovered extensive local processing and clustered plasticity of synaptic inputs that enhance the computational power of dendrites (Grienberger et al., 2015, Harvey and Svoboda, 2007, Kleindienst et al., 2011, London and Hausser, 2005, Losonczy et al., 2008). Although less studied, similar local processing appears to occur in terminal axon arbors, in which presynaptic inhibition and inhomogeneous distributions of voltage-gated ion channels can diversify the output of a single neuron (Debanne, 2004, Asari and Meister, 2012).

Amacrine cells (ACs) are a diverse class of interneurons in the retina (Helmstaedter et al., 2013, MacNeil and Masland, 1998). Most of the approximately 50 distinct AC types lack separate dendrites and axons, and, instead, receive input and provide output through the same neurites. The radially symmetric arbors of starburst ACs receive synaptic input and release neurotransmitters near and far from the soma, respectively (Ding et al., 2016, Vlasits et al., 2016). In a seminal study, Euler et al. (2002) discovered by two-photon Ca^2+^ imaging that the four to six primary neurites of starburst ACs with their daughter branches function as independent centrifugal motion sensors. Another AC type (A17) was recently shown to process converging inputs from rod bipolar cells separately (Grimes et al., 2010). For most AC types, however, whether neurites process inputs locally or integrate them globally, what stimulus features are encoded by neurites, and whether responses across neurite arbors are uniform or varied remains unknown.

VGluT3-expressing amacrine cells (VG3-ACs) stratify neurites broadly in the center of the inner plexiform layer (IPL). In somatic patch clamp recordings, VG3-ACs depolarize to light increments (ON) and decrements (OFF) constrained to a small area (i.e. receptive field center), but hyperpolarize to large ON and OFF stimuli that include the receptive field surround (Kim et al., 2015, Lee et al., 2014). VG3-ACs are dual transmitter neurons, which deploy their two transmitters in a target-specific manner. They provide glutamatergic input to W3 retinal ganglion cells (W3-RGCs) driving object motion sensitive responses (Kim et al., 2015, Krishnaswamy et al., 2015, Lee et al., 2014), and glycinergic input to Suppressed-by-Contrast RGCs (SbC-RGCs) inhibiting responses to small OFF stimuli (Lee et al., 2016, Tien et al., 2016, Tien et al., 2015). The target-specific use of excitatory and inhibitory transmitters (Tien et al., 2016, Lee et al., 2016), and the observation that removal of ON-OFF responsive VG3-ACs selectively affects OFF inhibition in SbC-RGCs (Tien et al., 2016) raise the question whether VG3-AC neurites process visual information locally or integrate it globally.

Here, we used two-photon Ca^2+^ imaging in a novel transgenic mouse line to analyze how stimulus contrast, size, and motion are processed in VG3-AC neurites.

## Results and discussion

We crossed *VG3-Cre* mice to a novel transgenic strain (*Ai148*) expressing the genetically encoded Ca^2+^ indicator GCaMP6f in a Cre-dependent manner enhanced by tTA-based transcriptional amplification. Staining for VGluT3 confirmed that GCaMP6f labeling in the inner nuclear layer (INL) and the IPL of *VG3-Cre Ai148* retinas was mostly restricted to VG3-ACs (Figure 1A-D) (Grimes et al., 2011, Kim et al., 2015). We imaged GCaMP6f signals in 23 x 23 μm scan fields in the IPL of flat-mounted retinas. Recording depths of scan fields were registered by their relative distance to the outer and inner boundaries of the IPL (0-100 %) detected by imaging transmitted laser light (Figure 1 – figure supplement 1). Visual stimulation (385 nm) was spectrally separated from GCaMP6f imaging (excitation: 940 nm, peak emission: 515 nm); and recordings were obtained from the ventral retina, where S-opsin dominates (Haverkamp et al., 2005, Wang et al., 2011). Ca^2+^ responses were no different for scanning at 9.5 Hz vs 37.9 Hz (Figure 1 – figure supplement 2). To maximize spatial resolution (pixel size: 0.29 x 0.36 μm), we therefore acquired all data in this study at 9.5 Hz. To identify processing domains in VG3-ACs neurites objectively, we automated detection of regions of interest (ROIs) in each scan field based using a serial clustering procedure (Figure 1E-H, s. Materials and methods).

**Figure 1.**
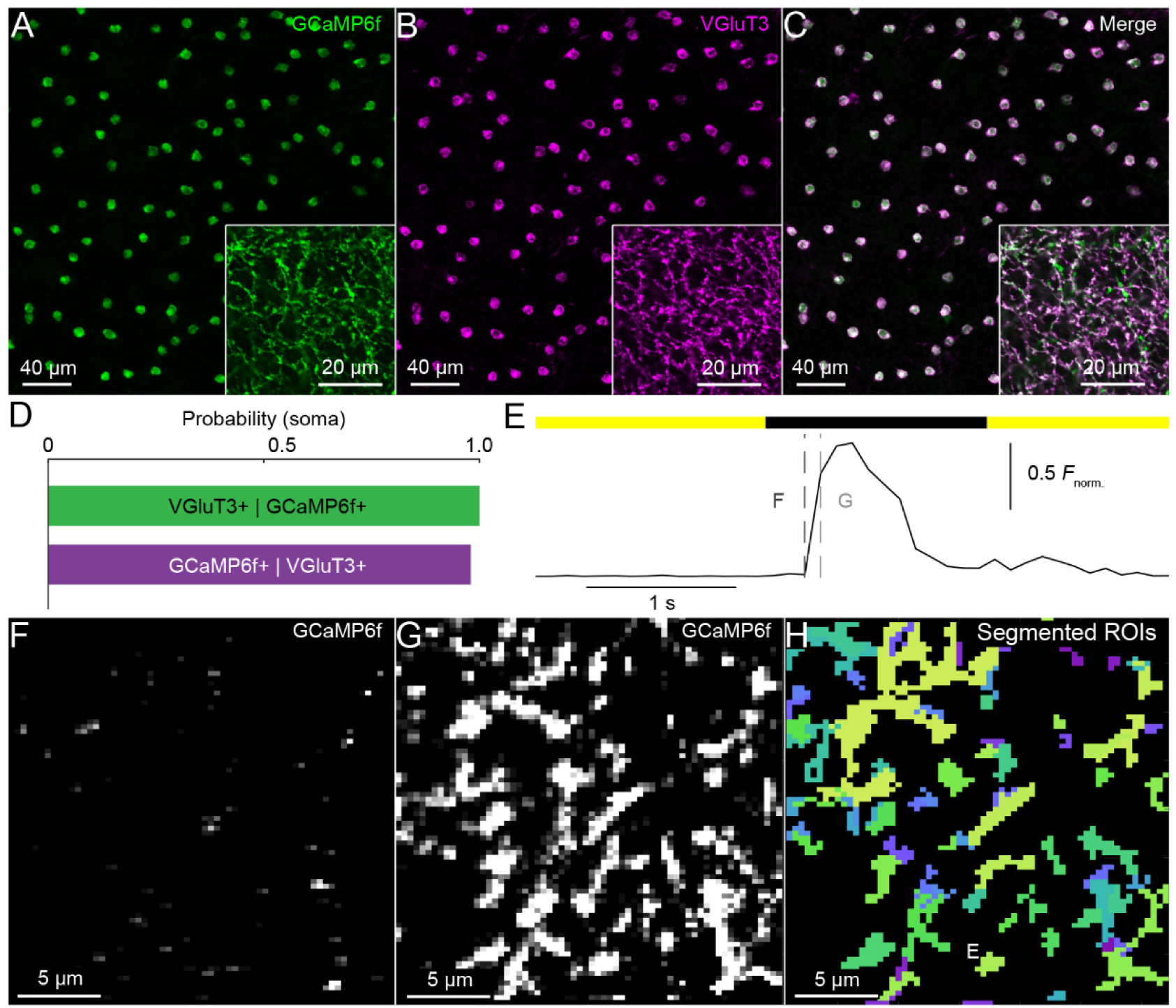
Specificity of GCaMP6f expression, VG3-AC neurite Ca^2+^ responses, and functional image segmentation. (**A-C**). Representative confocal images of the inner nuclear layer and IPL (insets) stained with anti-GFP, which recognizes GCaMP6f (**A, C**, green), and anti-VGluT3 (**B, C**, magenta). (**D**). Summary data (mean ± SEM) of the conditional co-localization probabilities of both signals. The green bar shows the probability that a GCaMP6f-positive cell (n = 111) is also VGluT3 positive. The purple bar shows the probability that a VGluT3-positive cell (n = 113) is also GCaMP6f positive. (**E**) A representative GCaMP6f response trace of an ROI marked in (**H**), responding to a small spot of light (radius: 50 μm). (**F-G**) Frames of a two-photon image series at time points indicated by dashed lines in (**E**). (**H**) Image segmentation of the scan field shown in (**F**) and (**G**) by a serial clustering procedure (see Materials and methods).

In somatic patch clamp recordings, VG3-ACs depolarize to small ON and OFF stimuli (Lee et al., 2014, Kim et al., 2015, Grimes et al., 2011). Somatic Ca^2+^ transients exhibited similar ON-OFF profiles to those observed in voltage recordings (Figure 2A,B). To test, whether these responses are uniformly distributed across VG3-AC neurites or not, we recorded Ca^2+^ transients elicited by contrast steps in a small spot (radius: 50 μm) at different depths of the IPL (Figure 2A,B and Video 1). We quantified contrast preference by a polarity index, ranging from -1 for pure OFF responses to 1 for pure ON responses (s. Materials and methods). Polarity indices varied widely between ROIs (n = 1120) (Figure 2C) and were correlated with IPL depth (R^2^ = 0.129, p < 10^-67^), as neurites in the outer IPL (depths < 40 %) responded more strongly to OFF stimuli, and neurites in the inner IPL (depths > 40 %) responded more strongly to ON stimuli (Figure 2D). Both ON and OFF responses were transient (Figure 2E); and in the inner IPL (depths > 40 %), ON responses were more transient than OFF responses (Figure 2F). Thus, VG3-AC neurites process visual information locally, not globally, and exhibit diverse contrast preferences and response kinetics in different layers of the IPL.

**Figure 2.**
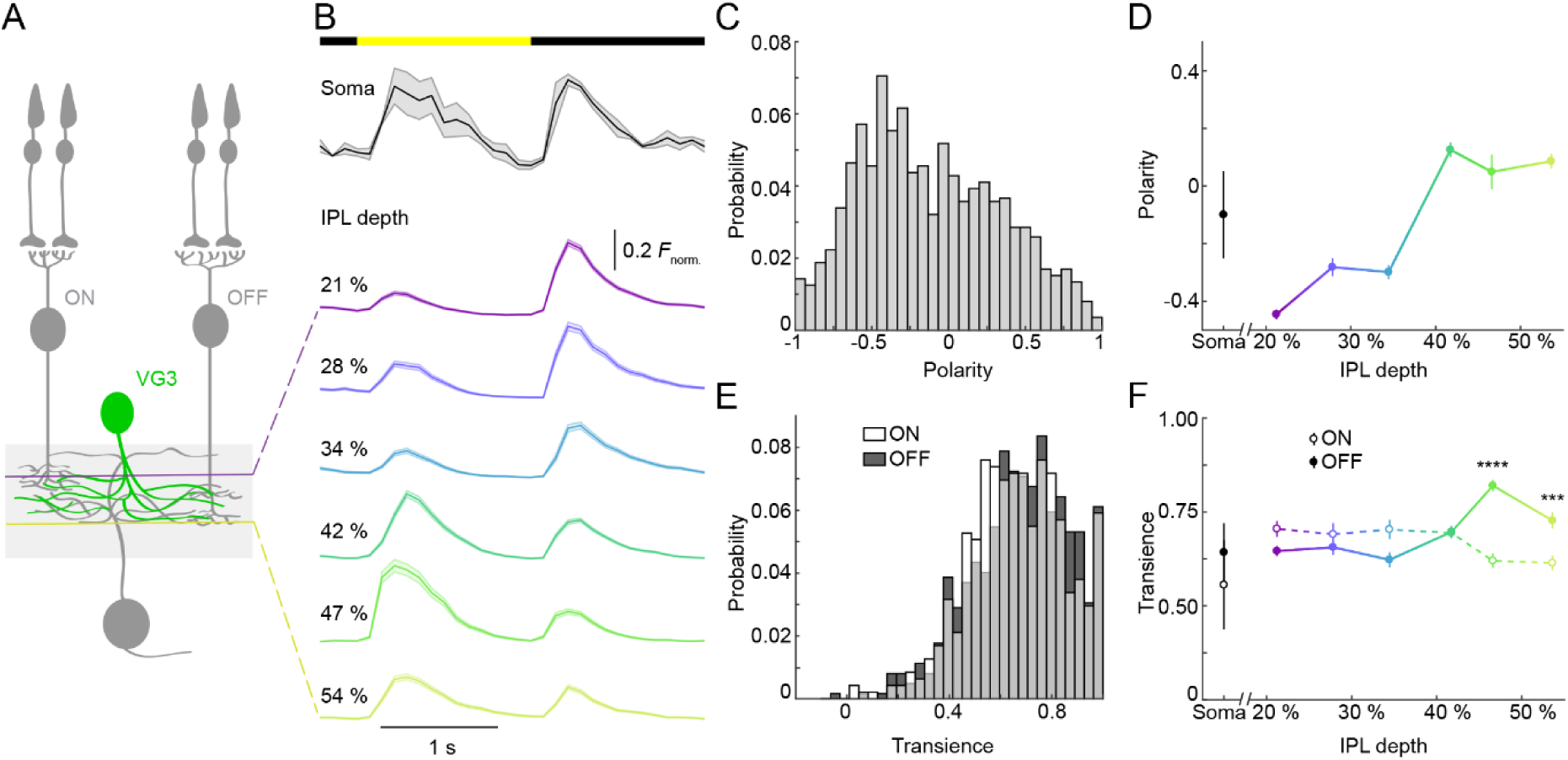
Responses of VG3-AC neurites to contrast steps vary across IPL depths. (**A**) Schematic of the VG3-AC circuit. VG3-AC neurites receive input from ON and OFF bipolar cells and synapse onto retinal ganglion cells. (**B**) Ca^2+^ transients of ROIs at different imaging depth to a contrast steps in a small spot (radius: 50 μm). A bar at the top indicates the stimulus timing. The black trace (shaded area) show the mean (± SEM) responses of VG3-AC somata. The six different color-coded traces (shaded areas) indicate the mean (± SEM) responses of ROIs at different IPL depths (21%: n = 249, purple; 28%: n = 120, blue; 34%: n = 143, sky; 42%: n = 248, green; 47%: n = 109, lime; 54%: n = 251, olive). (**C**) The distribution of polarity indices of VG3-AC neurite ROIs. (**D**) Summary data (mean ± SEM) of polarity indices as a function of IPL depth. Polarity indices differed between different IPL depths (p < 10^-65^, Kruskal-Wallis one-way ANOVA). ROIs at 21% IPL depth were more polarized toward OFF than at other depths (p < 0.01 compared to 28% and 34%; p < 10^-7^ for 42% - 54%). ROIs from 42% - 54% IPL depth were more polarized toward ON than ROIs from 21% - 34% (p < 10^-5^). (**E**) The distribution of transience indices of ROIs for ON (white) and OFF (dark gray) responses. (**F**) Summary data (mean ± SEM) of transience indices of ON (open circle) and OFF (filled circle) responses as a function of IPL depth. The transience of ROIs did not different significantly across IPL depth (p = 0.1364, main effect of depth, two-way ANOVA) or between ON and OFF contrast (p = 0.0695, main effect of contrast), but are differed in the interaction of depth and contrast (p < 10^-10^). For ROIs at 47% and 54% IPL depth, ON responses were more transient than OFF responses (p < 10^-5^ and p < 10^-3^, respectively).

A hallmark of VG3-ACs’ somatic voltage responses is strong size selectivity. We explored how VG3-AC neurites at different depths in the IPL respond to contrast steps in spots of different sizes. At all depths, only small stimuli (radius: < 200 μm) elicited Ca^2+^ transients (Figure 3A and Video 1). OFF responses preferred smaller stimuli than ON responses across most of the IPL (Figure 3B,C). Interestingly, variations in response center size between ROIs were correlated for ON and OFF responses (Figure 3 – figure supplement 1). Thus, stimulus size, like contrast, is encoded locally in VG3-AC neurites; and some mechanisms that shape size selectivity appear to be shared between ON and OFF responses of a given VG3-AC neurite.

**Figure 3.**
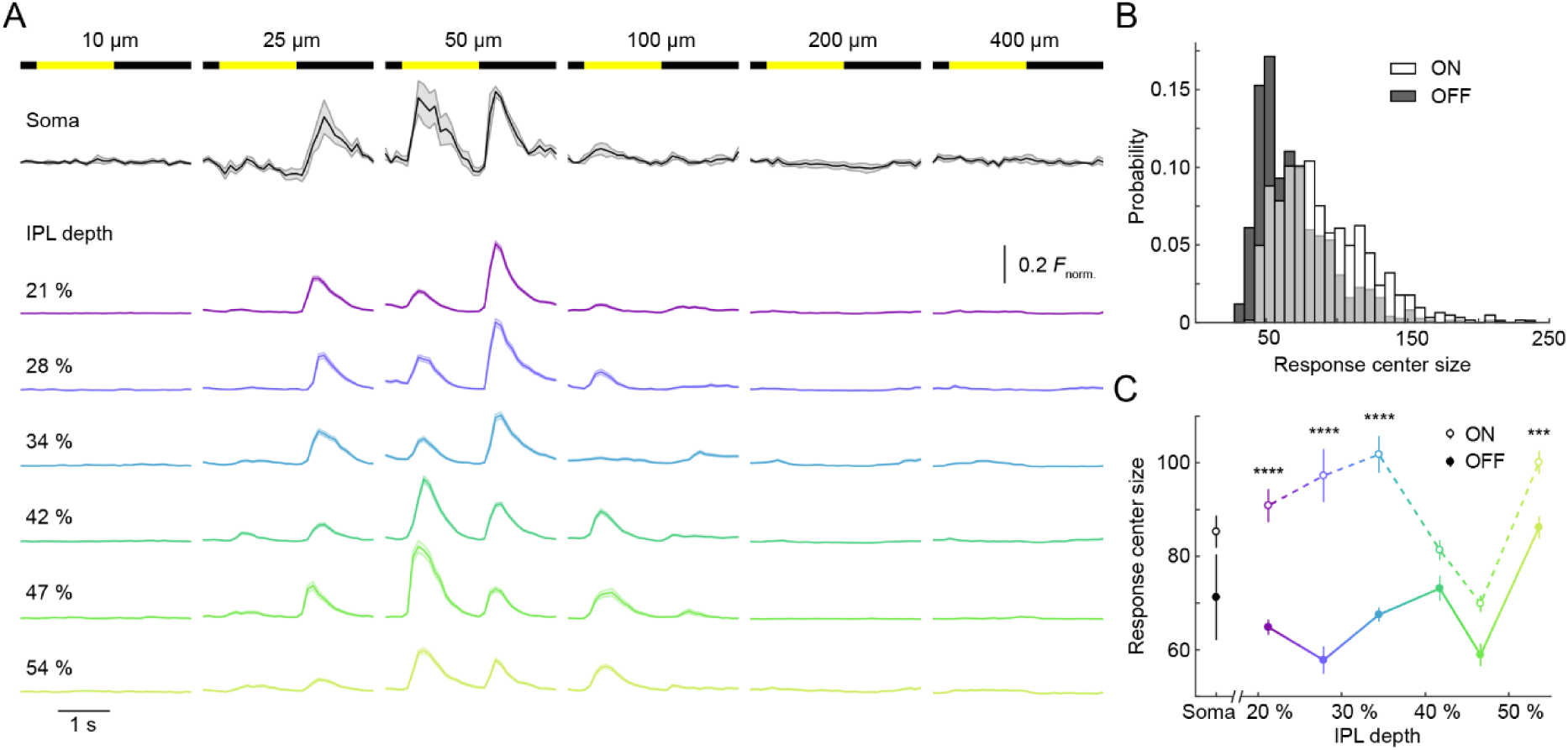
Size selectivity of ON and OFF responses in VG3-AC neurites. (**A**) Ca^2+^ transients of ROIs at different imaging depth to a contrast steps in spots of different size. Spot radii are noted at the top of bars indicating stimulus timing. The black traces (shaded areas) show the mean (± SEM) responses of VG3-AC somata. The six different color-coded traces (shaded areas) indicate the mean (± SEM) responses of ROIs at different IPL depths (21%: n = 249, purple; 28%: n = 120, blue; 34%: n = 143, sky; 42%: n = 248, green; 47%: n = 109, lime; 54%: n = 251, olive). (**B**) The distribution of preferred stimulus sizes of ON (white) and OFF (dark gray) responses of ROIs. (**C**) Summary data (mean ± SEM) of preferred stimulus sizes of ON (open circle) and OFF (filled circle) responses as a function of IPL depth. Preferred stimulus sizes differed between IPL depths (p < 10^-10^, main effect of contrast, two-way ANOVA) and between ON and OFF responses overall (p < 10^-10^, main effect of contrast, two-way ANOVA). Moreover, preferred stimulus sizes differed between ON and OFF responses of ROIs at all IPL depths (p < 10^-6^ from 21% - 34%, p < 0.001 at 54%) except for 42% (p = 0.21) and 47% (p = 0.59). The interaction between depth and contrast was also significant (p < 10^-8^).

VG3-ACs participate in object motion sensitive circuits in the retina, amplifying selectively the responses of W3-RGCs to local image motion (Krishnaswamy et al., 2015, Kim et al., 2015). We tested the ability of individual VG3-AC neurites to distinguish local and global image motion, using a stimulus in which square wave gratings overlaying center and surround regions of neurite receptive fields moved separately or together (Kim et al., 2015, Olveczky et al., 2003, Zhang et al., 2012). Isolated motion of the center grating elicited robust Ca^2+^ transients in VG3-AC neurites at all depths, which remained silent during simultaneous motion of gratings in center and surround (i.e. global motion) (Figure 4A and Video 2). As a result, local motion preference indices (s. Materials and methods) of > 83% of ROIs were > 0.9 (Figure 4B,C). Thus, contrary to the diversity of responses to contrast steps, VG3-AC neurites exhibit uniform object motion selectivity.

**Figure 4.**
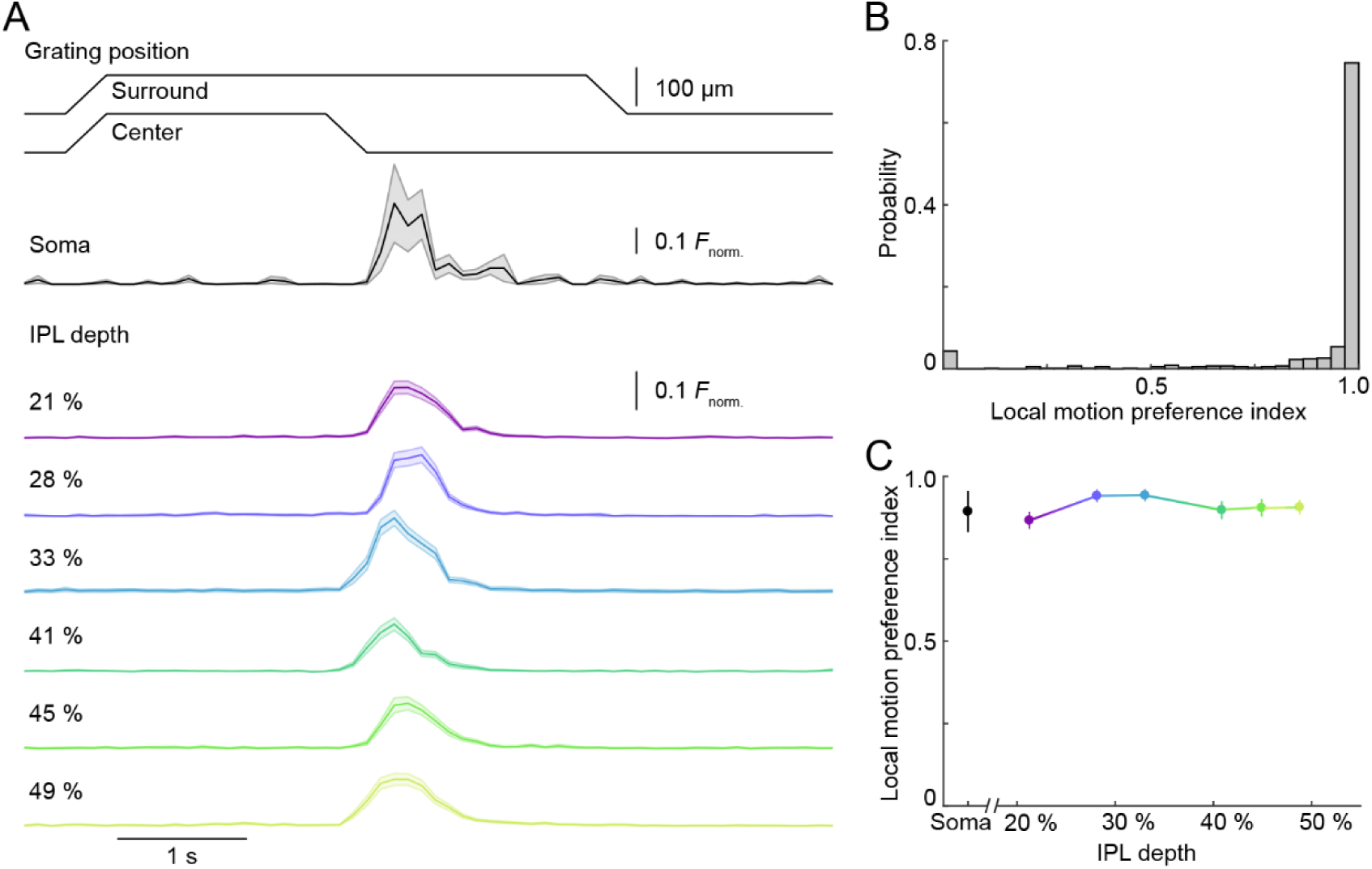
Uniform local motion preference of VG3-AC neurites. (**A**) Schematic at the top shows the time course of the grating motion in the receptive field center and surround (see Video 2 and Materials and methods). The black trace (shaded area) shows the mean (± SEM) responses of VG3-AC somata. The six different color-coded traces (shaded areas) indicate the mean (± SEM) responses of ROIs at different IPL depths (21%: n = 115, purple; 28%: n = 88, blue; 33%: n = 75, sky; 41%: n = 92, green; 45%: n = 89, lime; 49%: n = 116, olive). (**B**) The distribution of local motion preference indices of all ROIs. (**C**) Summary data (mean ± SEM) of local motion preference indices as a function of IPL depth. Local motion preference indices did not differ across IPL depths (p = 0.4427, Kruskal-Wallis one-way ANOVA). No ROI group at any depth was significantly different from any ROI group at another depth.

Our results show that VG3-ACs process visual information locally and exhibit diverse preferences for stimulus contrast and size across their neurite arbors. Changes in contrast preference with increasing IPL depth likely reflect a shift in input from OFF to ON bipolar cells and align with the stratification patterns of bipolar cell axons (Franke et al., 2017, Helmstaedter et al., 2013). The consistent differences in receptive field size of OFF and ON responses (Figure 3B) indicate separate origins (e.g. in OFF and ON bipolar cells, respectively), whereas correlated variations in ON and OFF receptive field sizes between neurites (Figure 3 – figure supplement 1) hint at a shared underlying mechanisms (e.g. varying strength of surround inhibition to VG3-AC neurites). In spite of their diverse responses to contrast steps, VG3-AC neurites show uniform object motion selectivity, which relies on rectified excitatory input from bipolar cells and strong surround inhibition from ACs (Kim et al., 2015). Local processing suggests that the output of VG3-ACs at different release site may convey different visual information. Whether release sites with different Ca^2+^ response profiles connect to different ganglion cell targets and/or use different neurotransmitters (glutamate vs. glycine) is an interesting question for future studies.

## Materials and methods

### Animals

We crossed VG3-Cre mice provided by Dr. R. H. Edwards (Grimes et al., 2010) to the *Ai148* strain, a novel transgenic line made by first targeting a Flp/Frt-based docking site cassette into the TIGRE locus on chromosome 9, followed by modification of that locus by Flp-induced RMCE. Ai148 mice contain Cre-regulated units within the TIGRE locus (Madisen et al., 2015) for both GCaMP6f and tTA2 expression, thereby allowing for tTA-based transcriptional amplification of GCaMP6f in a two mouse system. Mice were housed in a 12 hr light/dark cycle and fed *ad libidum*. We isolated retinas from mice of both sexes aged between postnatal day 30 (P30) and P45. All procedures in this study were approved by the Institutional Animal Care and Use Committee of Washington University School of Medicine (Protocol # 20140095) and were performed in compliance with the National Institutes of Health *Guide for the Care and Use of Laboratory Animals*.

### Tissue preparation

Mice were dark-adapted for more than 1 hour, deeply anesthetized with CO_2_, killed by cervical dislocation, and enucleated. Retinas were isolated under infrared illumination in mouse artificial cerebrospinal fluid buffered with HEPES (mACSF_HEPES_ for immunohistochemistry) or sodium bicarbonate (mACSF_NaHCO3_ for two-photon imaging). mACSF_HEPES_ contained (in mM): 119 NaCl, 2.5 KCl, 2.5 CaCl_2_, 1.3 MgCl_2_, 1 NaH_2_PO_4_, 11 glucose and 20 HEPES (pH adjusted to 7.37 with NaOH). mACSF_NaHCO3_ contained (in mM) 125 NaCl, 2.5 KCl, 1 MgCl_2_, 1.25 NaH_2_PO_4_, 2 CaCl_2_, 20 glucose, 26 NaHCO_3_ and 0.5 L-Glutamine equilibrated with 95% O_2_ / 5% CO_2_. Isolated retinas were flat mounted on black membrane disks (HABGO1300, Millipore for immunohistochemistry) or transparent membrane discs (Anodisc 13, Whatman, for two-photon imaging).

### Immunohistochemistry

Flat-mounted retinas were fixed for 30 min in 4% paraformaldehyde in mACSF_HEPES_ at room temperature (RT) and washed three times for 10 min in PBS at RT. The fixed tissue was cryoprotected with incubations in 10%, 20%, and 30% sucrose in PBS for 1 hr at RT, 1 hr at RT, and overnight at 4°C, respectively, followed by three cycles of freezing (held over liquid nitrogen) and thawing (in 30% sucrose in PBS). Retinas were then washed three times in PBS for 1 hr at RT, and stained for VGluT3 (rabbit anti-VGluT3, Cat. No. 1352503, Synaptic Systems) and GFP (chicken anti-GFP, 1:1000, Cat. No. A10262, ThermoFisher) for three to five days at 4°C in PBS with 5% normal donkey serum and 0.5% Triton X-100. Subsequently, retinas were washed three times for 1 hr in PBS, stained with Alexa 488- Alexa 568- conjugated secondary antibodies (Invitrogen, 1:1000) overnight at 4 °C, washed three times in PBS for 1 hr, and mounted in Vectashield mounting medium (Vector Laboratories) for confocal imaging.

### Confocal imaging

Confocal image stacks of fixed tissue were acquired through 20 X 0.85 NA or 60 X 1.35 NA oil immersion objectives (Olympus) on an upright microscope (FV1000, Olympus). Confocal images were processed and analyzed with Fiji (Schindelin et al., 2012).

### Visual stimulation

Visual stimuli were written in MATLAB (The Mathworks) using the Cogent Graphics toolbox (John Romaya, Laboratory of Neurobiology at the Wellcome Department of Imaging Neuroscience, University College London). Stimuli were presented from a UV E4500 MKII PLUS II projector illuminated by a 385 nm LED (EKB Technologies) and focused onto the photoreceptors of the ventral retina via a substage condenser of an upright two-photon microscope (Scientifica). All stimuli were centered on the two-photon scan field and their average intensity was kept constant at ∼ 1,600 S-opsin isomerizations / S-cone /s. To measure the contrast and size preference, the intensity of spots of varying radii (10, 25, 50, 100, 200, 300, and 400 μm) was square-wave-modulated (1.5 s ON, 1.5 s OFF) for 5 cycles. The order in which spots of different size were presented was randomly chosen for each scan field. To test responses to local *vs*. global motion stimuli, narrow square wave gratings (bar width: 50 μm) over the receptive field center (radius: 75 μm) and surround (150-800 μm from center of the image) were moved separately or in unison (Kim et al., 2015, Zhang et al., 2012). A gray annulus was included in the spatial layout of the stimulus to reliably separate movement in the center and surround. Each grating motion lasted 0.5 s, and movements were separated by 1.5 s.

### Two-photon imaging

A custom-built upright two-photon microscope (Scientifica) controlled by the Scanimage r3.8 MATLAB toolbox was used in this study; and images were acquired via a DAQ NI PCI6110 data acquisition board (National Instruments). GCaMP6f was excited with a Mai-Tai laser (Spectra-Physics) tuned to 940 nm, and fluorescence emission was collected via a 60 X 1.0 NA water immersion objective (Olympus) filtered through consecutive 450 nm long-pass (Thorlabs) and 513-528 nm band-pass filters (Chroma). This blocked visual stimulus light (peak: 385 nm) from reaching the PMT. Because we observed no qualitative differences between Ca^2+^ responses scanned at 9.5 Hz and 37.9 Hz (Figure 1 – figure supplement 2), we acquired 23 x 23 μm images (pixel size: 0.29 x 0.36 μm) throughout this study at 9.5 Hz. Imaging depths were registered by their relative distances to the borders between the IPL and the inner nuclear layer (0% IPL depth) and between the IPL and the ganglion cell layer (100% IPL depth). Borders were detected in transmitted light images (Figure 1 – figure supplement 1). Scan fields at different IPL depths were imaged in pseudorandom order; and for each scan the retina was allowed to adapt to the laser light for 30 s before presentation of visual stimuli. All images were acquired from the ventral retina, where S-opsin dominates (Wang et al., 2011). Throughout the experiments retina were perfused at ∼7 mL / min with 34°C mACSFNaHCO3 equilibrated with 95% O_2_ / 5% CO_2_.

### Image processing

#### Registration

Transmitted light images were acquired simultaneously with fluorescence images and were used to detect z-axis displacements that resulted in rejection of the respective image series. Images of series without z-axis displacements were registered to the middle frame using built-in functions in MATLAB. Rigid transformations were applied to both transmitted and fluorescence images. The quality of registration was confirmed by visual inspection, before transformed fluorescence images were used for further image processing and analysis.

#### Denoising

Time series of each pixel were searched for outliers (> 10 SD). If outliers were isolated in time (i.e. pixel value before and after outlier < 10 SD), they were replaced with the average of the value before and after the outlier. This algorithm effectively removed PMT shot noise.

#### Segmentation

To identify functional processing domains in VG3-AC neurites with minimal assumptions and user involvement, we developed a serial clustering procedure, in which a functional clustering algorithm is successively applied to different image features. This procedure removed pixels of the image not responding to visual stimulation and automatically assigned responsive pixels to functionally coherent, spatially contiguous regions of interest (ROIs). The functional clustering algorithm was based on Shekhar et al. (2016), beginning with principal components analysis to reduce the dimensionality of the input feature to the minimum needed to explain 80% of its variance. This was followed by a K-nearest-neighbor (KNN) algorithm, which generated a connectivity matrix. The connectivity matrix was then used in community detection clustering (Le Martelot and Hankin, 2012). We first applied functional clustering to the raw data of an image series and removed low-intensity pixels. Signals of remaining pixels were normalized to their peak and fed back into the functional clustering algorithm to group pixels with similar response properties. Groups of functionally similar pixels were divided into spatially contiguous ROIs within the image. The average response traces of these ROIs were subjected to further rounds of functional clustering, in which spatially adjacent ROIs that were grouped in the same cluster were merged. This process was repeated until it converged on a stable solution (typically less than 15 iterations). Finally, ROIs identified in this procedure were examined for signal correlation with the visual stimulus and size, to reject non-responsive and/or small (< 5 pixels) ROIs.

### Image analysis

#### Polarity index

Responses of each ROI to contrast steps in small spots (radius: 50 μm) were divided into ON and OFF periods (1.5 s each). The median peak response to 5 stimulus repeats during each periods was then used to calculate a polarity index as follows:

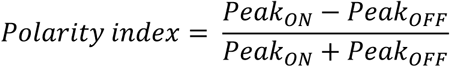

A polarity index of 1 indicates pure ON responses, whereas a polarity index of -1 indicates pure OFF responses.

#### Transience index

The transience index was calculated separately for ON and OFF responses of each ROI to contrast steps in small spots (radius: 50 μm) according to:

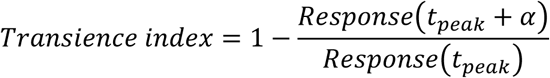

ON and OFF periods each lasted 1.5 s. *t_peak_* is the time to peak, measured from stimulus onset, and *α* is a delay set to the fourth frame (∼420 ms) after the peak frame. Transience was only calculated for ON and OFF responses that exceeded 25 % of the peak amplitude of the respective ROI. The maximum value of the transience index is 1, indicating that the GCaMP6f signal returned to baseline at time *α* after the peak.

#### Response center size

Because difference of Gaussians fits were poorly constrained by our data, we quantified stimulus size preference using a single nonparametric measure, the response center size. ON and OFF responses of each ROI were analyzed separately and their amplitudes plotted as a function of stimulus size. If responses to any size exceeded 25 *%* of the peak amplitude of the respective ROI, response center size was calculated as the center of mass of the stimulus size – response relationship.

#### Local motion preference index

Median responses of each ROI to isolated grating motion in the receptive field center (i.e. local motion) and to synchronous grating motion in receptive field center and surround (i.e. global motion) were used to calculate a local motion preference index as follows:

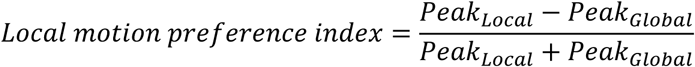

A local motion preference index of 1 indicates that the respective ROI responded only to local and not to global motion.

### Statistics

Functional imaging data were obtained from retinas of five mice. All summary data and response traces are presented as mean ± SEM. Differences between groups were statistically examined by two-way analysis of variance (for transience and response center size) with post hoc comparisons applying Student’s t-tests with Turkey’s HSD (honest significant difference) multiple comparison correction or by Kruskal-Wallis test (for polarity and local motion preference).

## Author contributions

The study was conceived and designed by J.-C.H. and D.K.; data were acquired by J.-C.H. and K.J.; data were analyzed and interpreted by J.-C.H. and D.K.; L.M. and H.Z. contributed unpublished essential reagents; and the manuscript was written by J.-C.H. and D.K. with input from all authors.

## Acknowledgments

We thank members of the Kerschensteiner lab for helpful comments and suggestions throughout this study. This work was supported by the National Institutes of Health (EY023341, EY026978, and EY027411 to DK and the Vision Core Grant EY0268) and by an unrestricted grant from the Research to Prevent Blindness Foundation to the Department of Ophthalmology and Visual Sciences at Washington University.

---

**Video 1. Ca^2+^ imaging of VG3-AC neurite responses to contrast steps in spots of varying size recorded at different IPL depths.**
Image series of GCaMP6f responses at 24% (middle) and 53% (right) IPL depth to contrast steps in spots of different size (left). The video is sped up 2.5-fold relative to the image acquisition. In the left panel, the area of the scan fields is indicated by a red box. Two average normalized ROI traces are shown at the bottom of the middle and the right panel.

**Video 2. Ca^2+^ imaging of VG3-AC neurite responses to motion stimuli recorded at different IPL depths.**
Image series of GCaMP6f responses at 24% (middle) and 49% (right) IPL depth to synchronous or isolated motion of square wave gratings in the center and surround separated by a gray annulus (left). The video is sped up 1.25-fold relative to the image acquisition. In the left panel, the area of the scan fields is indicated by a red box. Two average normalized ROI traces are shown at the bottom of the middle and the right panel.

**Figure 1 – figure supplement 1.**
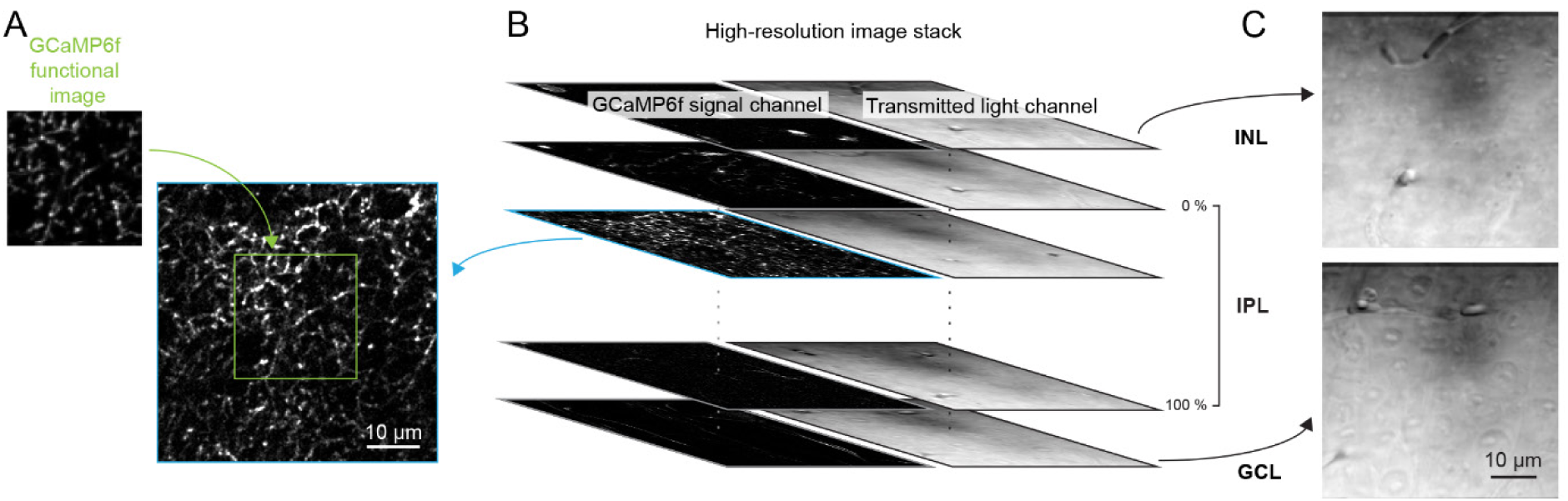
Registration of scan fields of functional GCaMP6f imaging to high-resolution image stacks to identify IPL depth. (**A, B**) Each functional imaging scan field of VG3-AC neurites (A, green, 64 x 80 pixels over 23 x 23 μm) was registered to one frame of a high-resolution image stack (B, blue, 512 x 512 pixels over 52.5 x 52.5 μm, 0.2 μm / z-step) acquired at the end of the functional imaging series. (**C**) Transmitted laser light was collected during acquisition of the high-resolution stack and used to identify the boundaries of the IPL. Top: transmitted light image of the inner plexiform layer (INL). Bottom: transmitted light image of the ganglion cell layer (GCL).

**Figure 1 – figure supplement 2.**
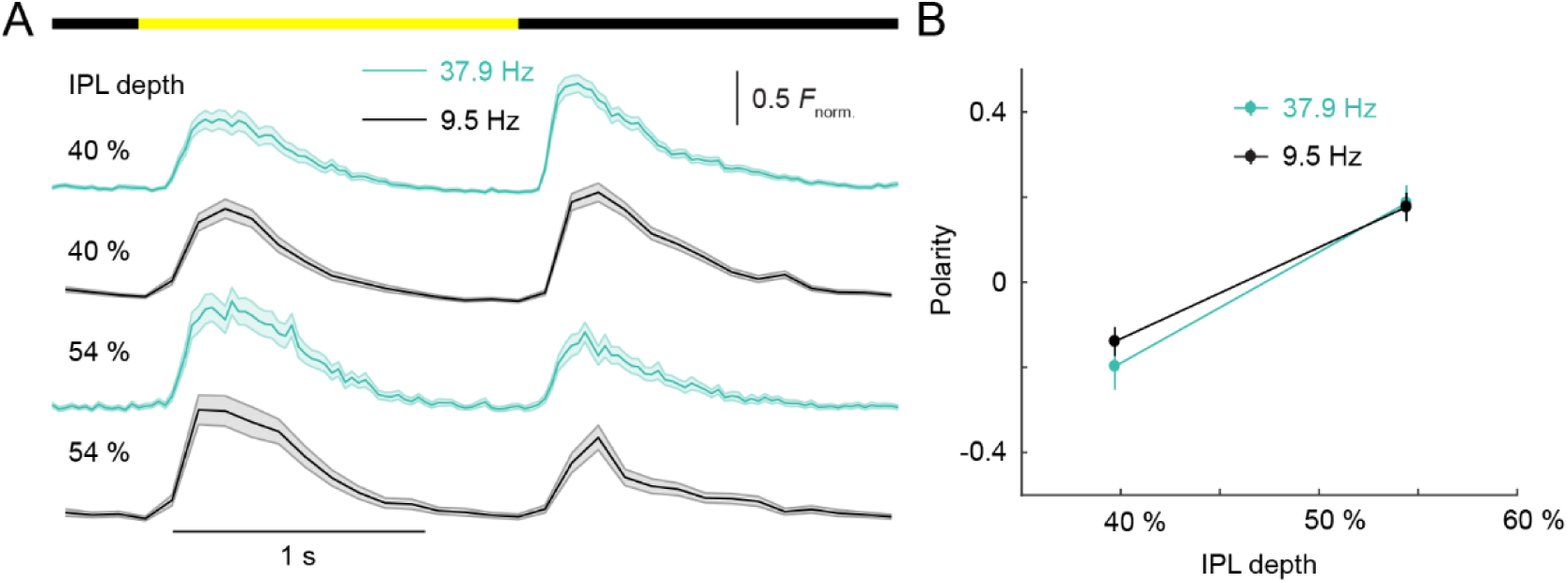
VG3-AC neurite responses did not differ between slower (9.5 Hz) and faster (37.9 Hz) scan rates. (**A**) Ca^2+^ transients of ROIs recorded at different scan rates from two different IPL depths. The bar at the top indicates the timing of stimulus. Black traces (shaded areas) show the mean (± SEM) responses of VG3-AC neurites scanned at 9.5 Hz; and green traces (shaded areas) show the mean (± SEM) responses of VG3-AC neurites scanned at 37.9 Hz. (40% at 9.5 Hz: n = 126; 40% at37.9 Hz: n = 67; 54% at 9.5 Hz: n = 62; 54% at 37.9 Hz: n = 28.) (**B**) Summary data (mean ± SEM) of polarity indices as a function of IPL depth for responses scanned at 9.5 Hz (black) and 37.9 Hz (green). The polarity of ROIs was different between IPL depths (p < 10^-10^, main effect of depth, two-way ANOVA), but did not differ significantly between scan rates (p = 0.62, main effect of frequency). The interaction between frequency and depth was not significant (p = 0.50).

**Figure 3 – figure supplement 1.**
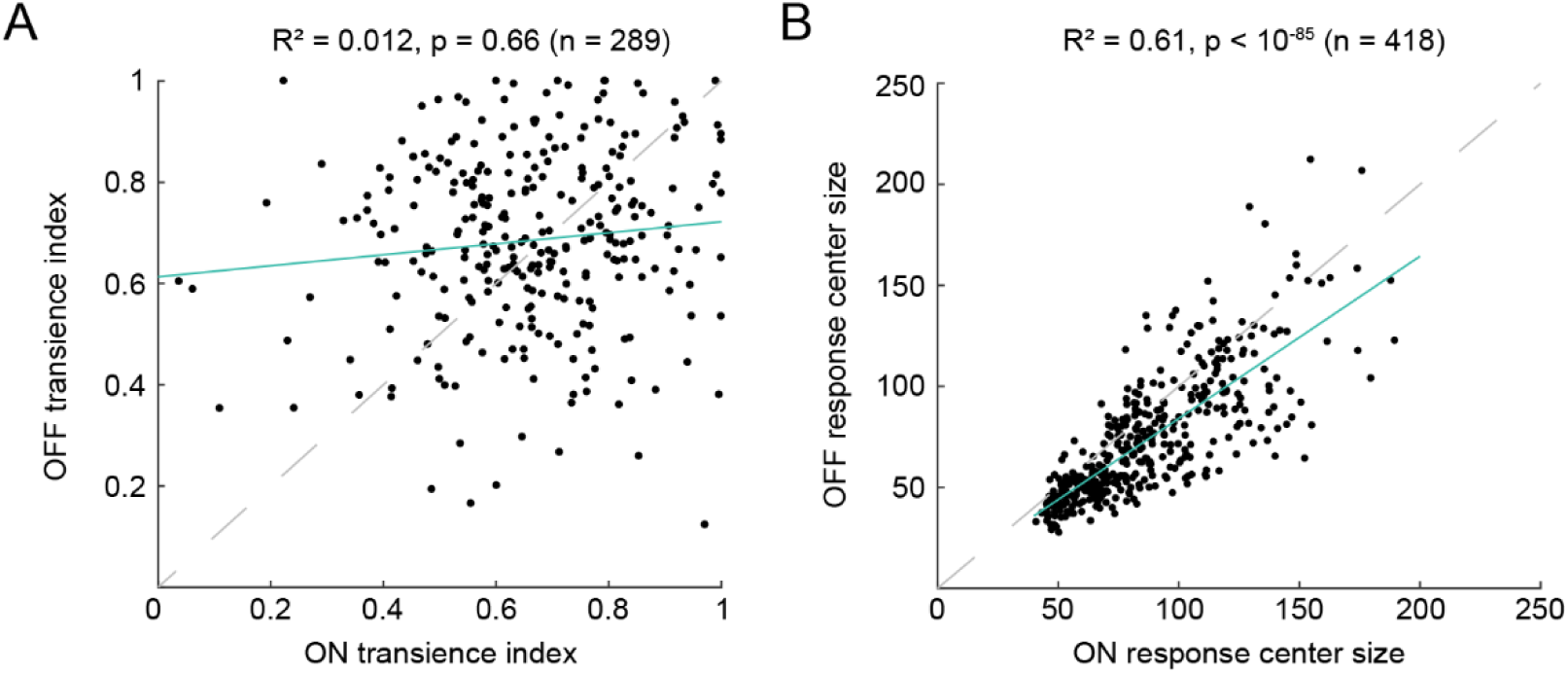
ON and OFF response center sizes, but not transience indices, of ROIs are highly correlated. (**A**) Correlation between ON and OFF transience indices was not significant (R^2^ = 0.012, p = 0.66, n = 289). If either ON or OFF response to 50-μm spots did not reach 25% of the maximal response of a given ROI, it was rejected from the analysis. (**B**) Correlation between ON and OFF response center size was significant (R^2^ = 0.61, p < 10^-85^, n = 418). If none of the ON or none of the OFF responses to spots of different sizes reached 25% of the maximal response of a given ROI, it was rejected from the analysis. Least square fits are shown as solid green lines and unity diagonals are indicated by dashed gray lines.

